# Tracking neural correlates of prioritizing working memory representations through retrospective attentional strengthening

**DOI:** 10.1101/2022.08.26.505307

**Authors:** Dongwei Li, Yiqing Hu, Mengdi Qi, Chenguang Zhao, Ole Jensen, Jing Huang, Yan Song

## Abstract

Previous work has proposed two potentials benefits of retrospective attention on working memory (WM): target strengthening and non-target inhibition. It remains unknown which hypothesis contributes to the improved WM performance, yet the neural mechanisms responsible for this attentional benefit are unclear. Here, we recorded electroencephalography (EEG) signals while 33 participants performed a retrospective-cue WM task. Multivariate pattern classification analysis revealed that only representations of target features were enhanced by valid retrospective attention during the retention, supporting the target strengthening hypothesis. Further univariate analysis found that mid-frontal theta inter-trial phase coherence (ITPC) and ERP components were modulated by valid retrospective attention and correlated with individual differences and moment-to-moment fluctuations on behavioral outcomes, suggesting that both trait- and state-level variability in attentional preparatory processes influence goal-directed behavior. Furthermore, task-irrelevant target spatial location could be decoded from EEG signals, indicating that enhanced spatial binding of target representation promotes high WM precision. Importantly, frontoparietal theta-alpha phase-amplitude-coupling was increased by valid retrospective attention and predicted the reduced randomly guessing rates. This long-range connection supported top-down information flow in engagement of frontoparietal networks, which might organize attentional states to integrate target features. Altogether, these results provide neurophysiological bases that retrospective attention improves WM precision through enhancing representation of target and emphasize the critical role of frontoparietal attentional network in the control of WM representations.

## Introduction

The cognitive resources of working memory (WM) are limited but flexible, in which attention acts as a gate, selectively allowing task-relevant information to enter WM, thereby supporting upcoming behaviors (1). Although rich evidence supported attentional promotion on WM representations, the underlying cognitive processing and neural substrates remain unclear. Additional, WM plays a fundamental role in fluid intelligence and broad cognitive function (2,3). Studying how attention benefits WM provides new insight into training individuals with memory deficits.

Previous work proposed two distinct hypotheses of attentional benefits on WM: target strengthening (4) and non-target inhibition (5). Several studies demonstrated the costs of invalid anticipatory attention (6,7), and larger behavioral benefits were observed as more non-targets were removed from WM with increasing memory loads (8). However, recent behavioral studies claimed that attending to the cued object in WM leads to stronger binding of that object to its context, without affecting the strength of non-cued objects (9). Extensive studies focused on behavioral performance (10), and even with several ERP/fMRI studies (11,12), it cannot be easy to distinguish different neural responses to targets and non-targets when they are presented simultaneously. Therefore, it is still lack of direct neurophysiological evidence to support one of the two distinct hypotheses of attention.

One way to examine the attentional effect on WM is to adopt retrospective-cues (retro-cue) under WM tasks (13,14). The pre-cue WM task has been widely used previously (15,16). In pre-cue tasks, cue is presented before the encoding period and reduces the memory load, confusing the retrieval of pure attentional effects. However, in retro-cue tasks, retro-cue is presented during the retention and obtains cleaner attentional effects. Previous studies found that retro-cue could modulate attention during retention (17) and prioritize relevant representations (18). Attention-related modulatory signals interacted with mnemonic representations (19), indicating the flexibility of information held in WM.

The posterior alpha activity (8 – 12 Hz), frontal theta activity (3 – 7 Hz) and their connections have been found to be related with attentional modulations. Abundant studies reported the relationship between visual attention and alpha event-related desynchronization (ERD; 20). Previous work has found that retro-cue modulated the alpha power in posterior areas during the retention (21,22), which supports that alpha oscillation acted as a gate to allow task-relevant information into attention (23,24). Mid-frontal theta activity was related to the attentional control (25) and played a vital role in the goal-related prioritization during WM (19,26). Previous studies found that midline theta power was involved in the control of WM (27), however, whether theta phase also contributes to the modulation in WM is unclear. Furthermore, connections between frontal and parietal cortex were related the top-down information flow (28,29). Previous work reported a correlation between frontal theta power and posterior alpha power (30). As such, the information flow in WM controlled by retrospective attention might be related with frontal theta parietal alpha connection.

Here, we adopted a visual WM precision task with valid and neutral retro-cues (**Fig 1A**) and recorded scalp electroencephalography (EEG) signals to directly examine the neurophysiological correlates of attentional effect on WM precision with both multivariate and univariate approaches. Multivariate decoding was applied to examine target enhancement (**Fig 1D**) and non-target inhibition hypothesis (**Fig 1E**). We further explored oscillatory correlates and connections to provide neurophysiological evidence that how retro-cue benefits WM representations.

**Fig 1.**
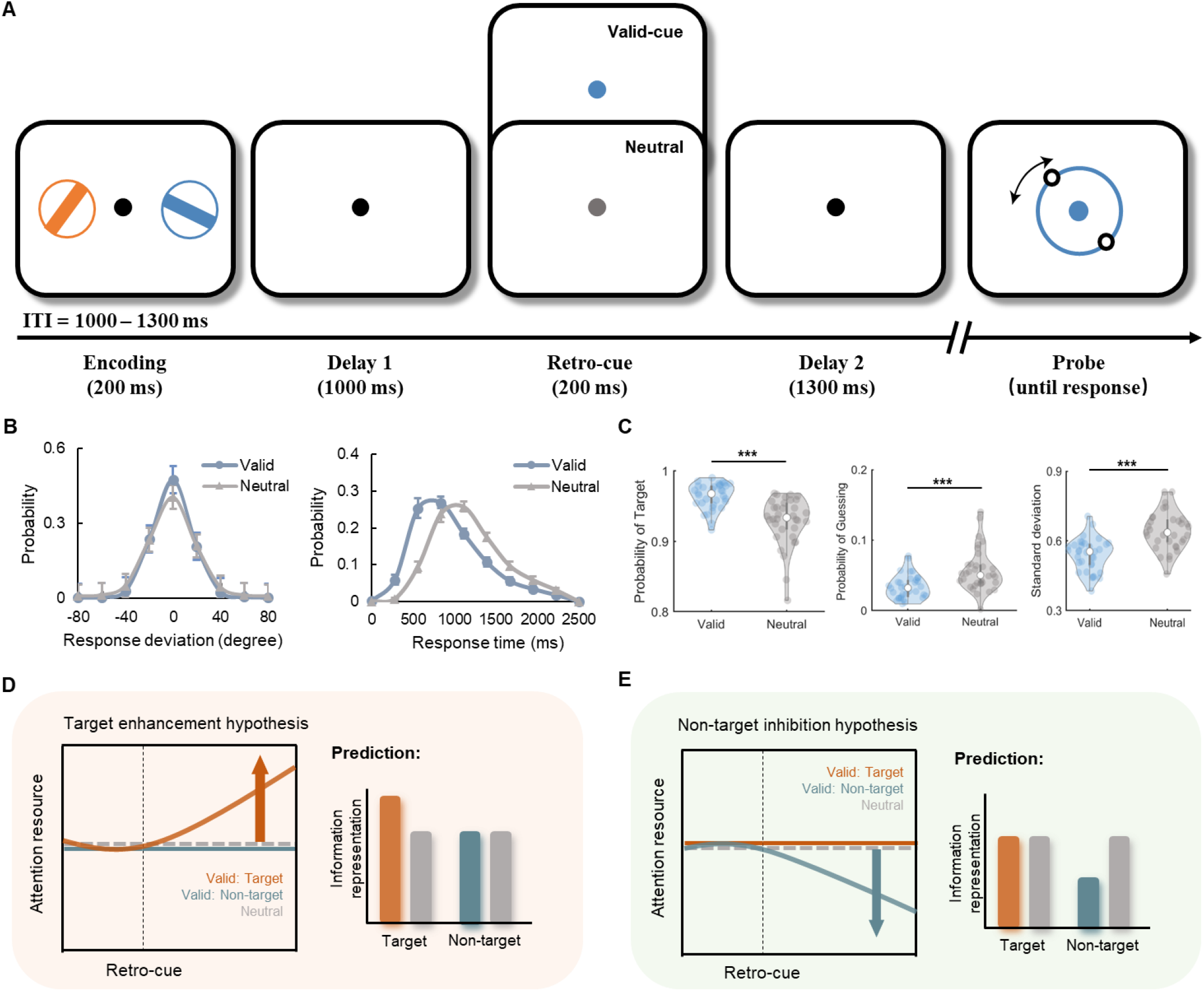
WM precision task and behavioral outcomes. **(A)** Trial sequences of the WM precision task with valid and neutral cues. **(B)** Probability of the response deviation and response time in both the valid and neutral conditions. **(C)** Mixture modeling of the behavioral results. **(D)** Target enhancement hypothesis. We predict a significant difference between valid retro-cue and neutral cue in the target orientation classification, but no difference between valid retro-cue and neutral cue in the non-target orientation classification. **(E)** Non-target inhibition hypothesis. We predict a significant difference between valid retro-cue and neutral cue in the non-target orientation classification, but no difference between valid retro-cue and neutral cue in the target orientation classification. ITI, inter-trial-interval; ^***^*p* < .001.

## Results

As illustrated in **Fig 1A**, participants were instructed to remember the exact orientations of the two colorful peripheral bars during the encoding period and to report the orientation of the target bar during the probe period with manipulating a mouse. Before the probe retrieval period, the color of the central black dot would change to the color of the target bar to retrospectively cue which one of the remembered two bars would be the target bar in advance in half of trials (valid condition), or change to gray giving no information about target in the other half of trials (neutral condition). Retro-cue related attentional benefits were evaluated by calculating behavioral and neural differences between valid and neutral conditions.

### Behavioral benefits from retro-cue

Robust behavioral benefits were found after valid retro-cues (**Fig 1BC**). Participants showed lower recall errors (*t*_*32*_ = 8.820, *p* < .001, *d* = 3.118) and faster reaction times (RTs; *t*_*32*_ = 18.192, *p* < .001, *d* = 6.432) in the valid condition. Then, maximum likelihood estimates were used to fit the response error distribution with the mixture model (See methods for more details). The modeling results also showed that valid retro-cues enhanced the probability of the target response (*t*_*32*_ = 6.444, *p* < .001, *d* = 2.278), improved the recall precision (*t*_*32*_ = 11.790, *p* < .001, *d* = 4.168), and reduced the probability of the randomly guessing (*t*_*32*_ = 4.379, *p* < .001, *d* = 1.548).

### Target and Non-target Orientation Decoding Support the Attentional Strengthening Hypothesis

To directly examine the information representations with two distinct hypotheses proposed in **Fig 1DE**, multivariate pattern classification was applied to train the EEG data with different labels to identify whether the target and non-target orientations could be decoded from neural response distributions (**Fig 2A**; see Methods). If the data support the target strengthening hypothesis, since the target representation is enhanced by valid retro-cue, we would expect a higher decoding accuracy of target orientation classification in valid condition, but no difference in decoding accuracy of non-target orientation classification. If the data support the non-target inhibition hypothesis, non-target representation will be inhibited by valid retro-cue. Since rejection template formation of distractor could achieve decoding accuracy of distractor information above chance level (31), we would expect a higher decoding accuracy of non-target orientation classification in valid condition, but no difference in decoding accuracy of target orientation classification.

**Fig 2.**
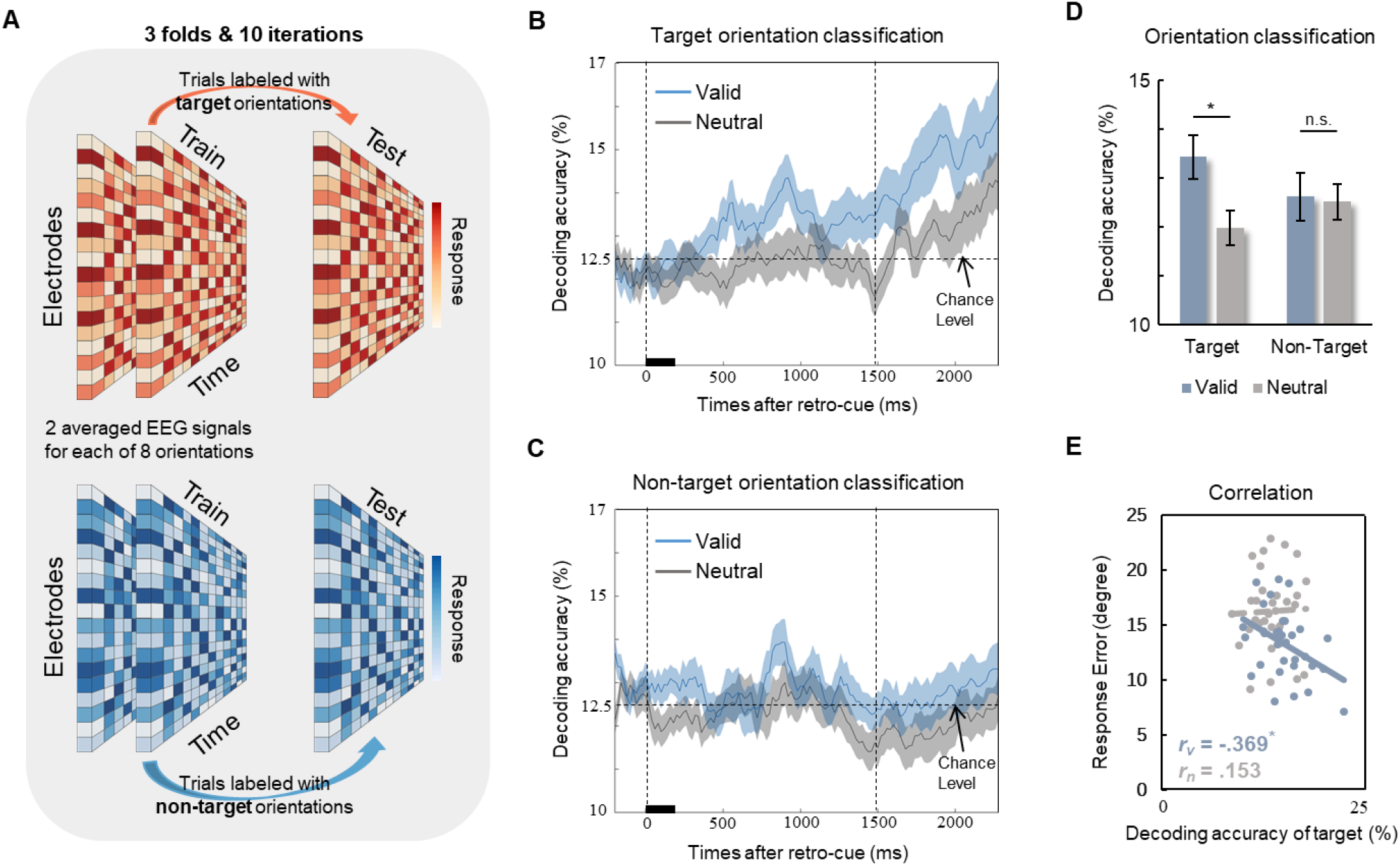
Orientation classifications. **(A)** Three-fold cross-validation was applied to trials labeled with target orientations (upper) and non-target orientations (below) to calculate the decoding accuracy of the multivariate pattern classification. **(B)** Temporal dynamics of the decoding accuracy for target orientation classification in the valid and neutral conditions. **(C)** Temporal dynamics of the decoding accuracy for non-target orientation classification in the valid and neutral conditions. **(D)** Averaged decoding accuracy of target and non-target orientation between 500 and 800 ms after the retro-cue supports the target enhancement hypothesis. **(E)** Individuals with higher decoding accuracy of the target orientation classification in the valid but not neutral conditions showed lower behavioral response errors. ^*^*p* < .050; n.s. represents ‘not significant’.

We found that valid retro-cues induced a higher decoding accuracy than neutral cues in target orientation classification (*t*_*32*_ = 2.388, *p* = .023, *d* = 0.844; **Fig 2B**), but no difference was found in non-target orientation classification (*t*_*32*_ = .086, *p* = .932, *d* = .030; **Fig 2C**), supporting the target strengthening hypothesis (**Fig 2D**). Importantly, increased decoding accuracy of the target orientation after valid retro-cues was correlated with reduced response errors (*r* = –.369, *p* = .035), but not after neutral retro-cues (*r* = .152, *p* = .397), indicating that enhanced target representation by valid retro-cue (**Fig 2E**).

### Flexible Temporal Dynamic in Color and Space Decoding

Then, we decoded the space and color of target to investigate whether features are binding to the goal-directed orientation to promote target representations. Both the color and space of the target could be decoded after retro-cue onset (**Fig 3AB**), suggesting that the priority of the target was modulated by object-based attention, not just orientation-based attention. A repeated-measures ANOVA was performed with 2 cue-types (valid, neutral) and 2 time periods (after retro-cue, after probe) as factors. Significant interactions were found in both the color (*F* = 23.896, *p* < .001, *η*_*p*_^*2*^ = .428) and space (*F* = 50.534, *p* < .001, *η*_*p*_^*2*^ = .612) classifications, indicating the different decoding patterns of the valid and neutral conditions between the time after the retro-cue and after the probe. These results suggested that once the spatial information was retrieved after the retro-cue, it was not necessary to decode this spatial representation again after the probe. More importantly, a significant negative correlation was found between space decoding accuracy after retro-cue and behavioral memory errors (*r* = –.355, *p* = .043; **Fig 3C**), indicating that effective spatial binding could benefit WM representational precision.

**Fig 3.**
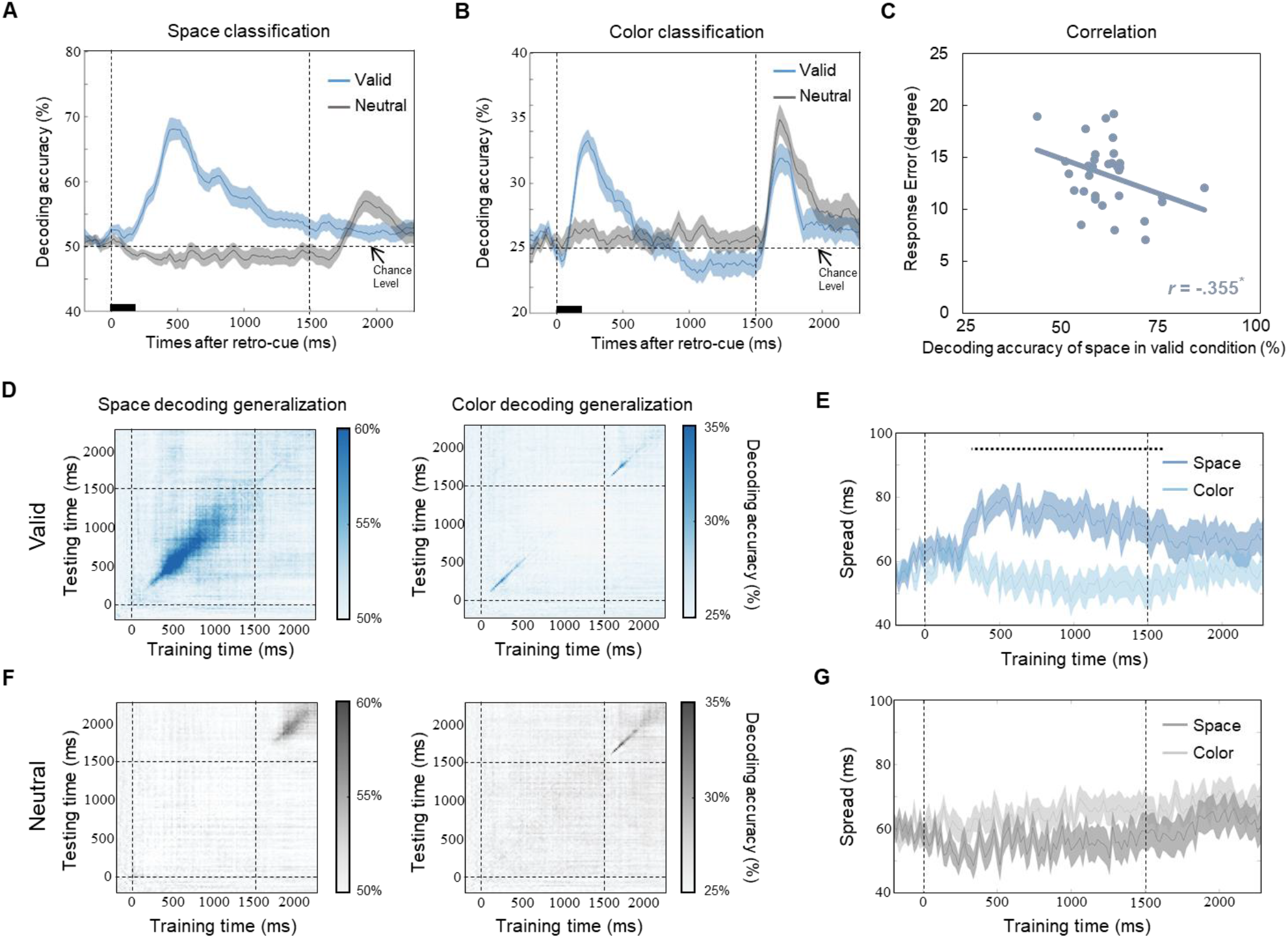
Target color and space classification. **(A)** Temporal dynamics of the decoding accuracy for target space classification in the valid and neutral conditions. **(B)** Temporal dynamics of the decoding accuracy for the target color classification in the valid and neutral conditions. **(C)** Higher decoding accuracy of space predicted the lower response error in valid condition. **(D)** Temporal generalization of the decoding accuracy for the space and color classification of the target in the valid condition. **(E)** Temporal dynamics of spreading time for target color and space decoding accuracy in valid condition. The black dotted line represents a time period with a significant difference. **(F)** Temporal generalization of the decoding accuracy for the space and color classification of the target in the neutral condition. **(G)** Temporal dynamics of spreading time for the target color and space decoding accuracy in the neutral condition. ^*^*p* < .050.

Temporal generalization analysis was conducted to investigate the representation dynamics of the different features (**Fig 3DF**). This analysis used representations at one time point to decode the same feature in another time points and a higher than chance level off-diagonal decoding accuracy means that neural representations remain stable across time (see Methods for more details). Space decoding generalization showed a larger temporal spread than color decoding generalization after valid retro-cues (*p*_*cluster*_ < .05; **Fig 3E**) but it did not show any difference after neutral cues (*p*_*cluster*_ > .05; **Fig 3G**), suggesting a stable representation of spatial information after valid retrospective attention.

### Frontal ERP and Theta Phase Track Attentional Control during WM

To investigate how neural activities contribute to retro-cue-related enhancement of target representations, we focused on the cue-evoked ERP component during the time with relatively high decoding accuracies of target orientation and spatial location. A large difference between valid and neutral condition during 600 to 1200 ms (N600) was found in frontal-central electrodes. The topographic map of N600 was illustrated in **Fig 4B**. Grand-average ERP waves across white dotes marked frontal electrodes (**Fig 4A**) showed a significant difference in averaged frontal N600 between valid and neutral conditions (*t*_*32*_ = 5.023, *p* < .001, *d* = 1.776). Importantly, a significant correlation (*r* = .400, *p* = .021; **Fig 4D**) was found between ERP retro-cue effect (difference in frontal N600 between valid and neutral conditions) and behavioral retro-cue effect (difference in recall errors between valid and neutral condition), indicating that larger difference in frontal N600 between two conditions predicted larger behavioral benefits on WM precision across individuals. A marginally significant correlation between ERP retro-cue effect and target orientation decoding accuracy in valid condition was further found (*r* = .336, *p* = .056), which might link the ERP component with decoding accuracy and partly explain how retrospective attention enhances representational precision in WM. Furthermore, when we divided trials into three equal bins according to the behavioral RTs (**Fig 4C**), a repeated-measures ANOVA on frontal N2 amplitudes showed a significant main effect on RT (*F* = 6.981, *p* < .001, *η*_*p*_^*2*^ = .179), indicating frontal N2 as a function of RTs across trials. These results suggested that retro-cue evoked frontal ERP could track both precision (N600) and speed (N2) benefits of WM behavioral outcomes.

**Fig 4.**
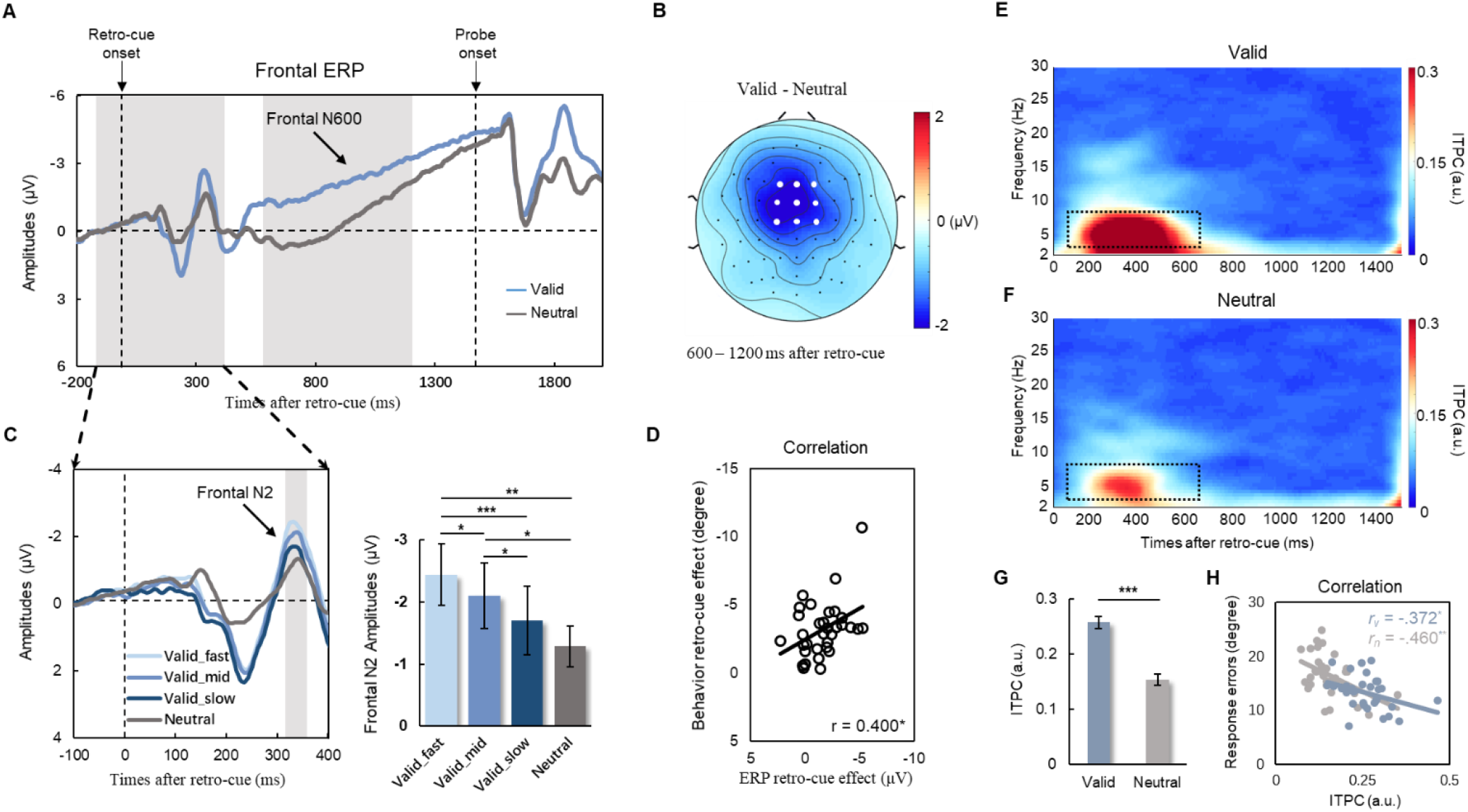
ERP and ITPC results. **(A)** Grand average waveforms of frontal ERPs for valid and neutral conditions. **(B)** An averaged topographic map (600 – 1200 ms) of the difference in frontal N600 amplitudes between valid and neutral conditions. White dots represented the electrodes used to calculate frontal N600. **(C)** Grand average waveforms of frontal ERPs for neutral condition and valid conditions with three different RTs. Frontal N2 amplitudes as a function of behavioral RTs across trials. **(D)** ERP retro-cue effect (frontal N600 in valid condition – frontal N600 in neutral condition) was correlated with behavioral retro-cue effect (recall errors in valid condition – recall errors in neutral condition). Time frequency representations of ITPC in both valid **(E)** and Neutral **(F)** conditions. **(G)** A higher theta ITPC averaged between 100 – 700 ms in midline frontal theta (3 – 7 Hz) was found in valid condition than in neutral condition. **(H)** Higher theta ITPCs predicted lower behavioral response errors in both valid and neutral conditions. ^*^*p* < .05; ^**^*p* < .01; ^***^*p* < .001.

To some extent, the ERP wave is related to phase-locked characteristic of oscillatory activities. Exploring the oscillatory mechanism of retro-cue benefits could help us understand the frequency contributions to abovementioned frontal ERP components (**Fig 4EF**). We found that inter-trial phase coherence (ITPC) in midline frontal theta (3 – 7 Hz) in valid condition was significantly higher than neutral condition (*t*_*32*_ = 11.687, *p* < .001, *d* = 4.132; **Fig 4G**), suggesting a better information integration and cognitive control in prefrontal cortex after valid retrospective attention. More importantly, significant correlations were found between theta ITPC and response errors in both valid (*r* = –.372, *p* = .033) and neutral (*r* = –.460, *p* = .007) conditions, indicating the important role of ITPC in target representation with stronger theta phase coherence leading to precise representation and lower behavioral response errors.

### Alpha Power and Long-range Connection within Frontoparietal Network

To examine how valid retro-cue modulated spatial attention to enhance target representation, we calculated ERD in alpha band (8 – 12 Hz) oscillatory activities during the retention of visual WM. As illustrated in **Fig 5AB**, a significant ERD in alpha power over parieto-occipital sensors was found after retro-cues in time frequency representations of both valid (*t*_*32*_ = –6.775, *p* < .001, *d* = 2.395) and neutral (*t*_*32*_ = –4.398, *p* < .001, *d* = 1.555) conditions. The alpha ERD was larger in valid condition than neutral condition (*t*_*32*_ = –5.606, *p* < .001, *d* = 1.982; **Fig 5CE**), and the corresponding topographic map of the difference in alpha power between valid and neutral conditions was shown in **Fig 5D**. This result suggested that more attention resources were deployed after the valid retro-cue during the retention.

**Fig 5.**
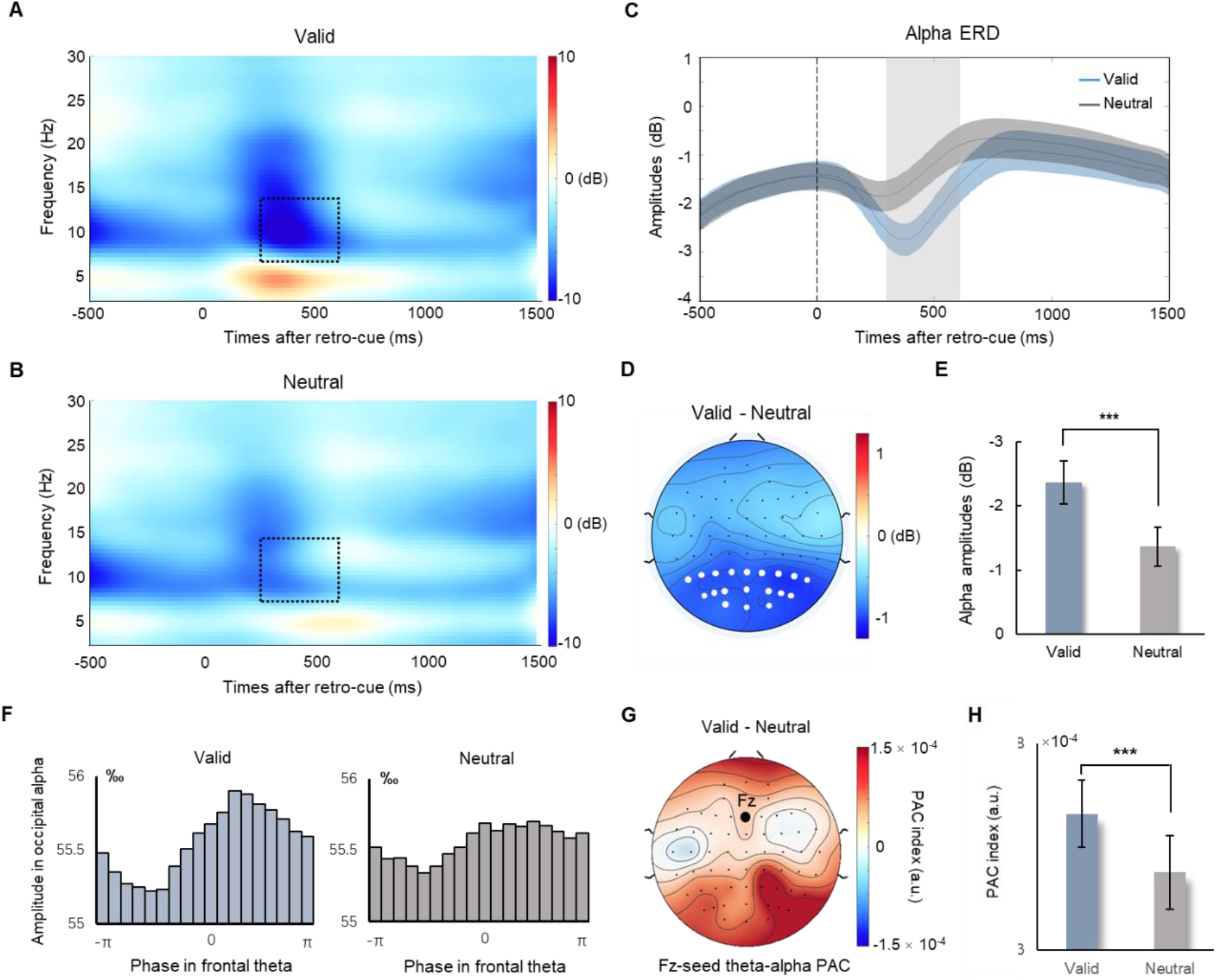
Oscillatory results. Time frequency representations in valid **(A)** and neutral **(B)** conditions. Black dashed squares represented alpha power (8 – 12 Hz) between 300 – 600 ms. **(C)** A larger alpha ERD was found in valid condition than in neutral condition. Grey area represented the duration (300 – 600 ms) used to calculate averaged alpha power. **(D)** An averaged topographic map (300 – 600 ms) of the difference in alpha power between valid and neutral conditions. White dots represented the electrodes used to calculate alpha power. **(E)** Averaged alpha power in valid and neutral conditions showed significant difference. **(F)** The alpha amplitudes across different theta phase bins in valid and neutral conditions. **(G)** A topographic map of the difference between valid and neutral conditions in theta-alpha PAC between Fz and all electrodes. **(H)** PAC index calculated between Fz and all parieto-occipital electrodes was larger in valid condition than in neutral condition. ^***^represented *p* < .001.

Since retro-cue could enhance target representations by modulating both frontal theta phase and posterior alpha power, we then investigated how these modulations were binding to the target by examining the frontal-posterior long-range connection through theta-alpha phase-amplitude-coupling (PAC; see Methods). The alpha amplitudes across different theta phase bins (**Fig 5F**) showed a curved uneven distribution in valid condition, but relatively even distribution in neutral condition. A topographic map of the difference between valid and neutral conditions in theta-alpha PAC between Fz and all electrodes was shown in **Fig 5G**. We found a stronger theta-alpha PAC between Fz and all parieto-occipital electrodes in valid condition than in neutral condition (*t*_*32*_ = 4.818, *p* < .001, *d* = 1.703; **Fig 5H**). Importantly, a significant correlation (*r* = –.372, *p* = .033) was found between Δtheta-alpha PAC (difference between valid condition and neutral condition) and Δprobability of guessing, suggesting that larger retro-cue effect in theta-alpha PAC predicted lower behavioral guessing rates. These results indicated that frontoparietal connection might be a fundamental mechanism of retrospective attention during WM retention.

## Discussion

How attention prioritizes relevant information held in WM to support flexibility of WM is the foundation of how brain resolve the limit of WM resource. In the present study, we applied both multivariate and univariate analysis to investigate whether and how retrospective attention improved WM performance with a typical retro-cue WM task. Only representations of target were enhanced and prioritized by retro-cue, which supports the target strengthening hypothesis. The task-irrelevant feature of target, spatial location, could be decoded with high priority in valid condition, indicating spatial information binding with the target object promotes WM representation precision. Furthermore, mid-frontal theta ITPC predicted individual differences of WM precision across participants and N2 components tracked moment-to-moment fluctuations of reaction time within participants. In addition, spatial attention-related parietal alpha ERD and its coupling with mid-frontal theta phase were modulated by valid retro-cue, further emphasizing neural correlates and connections within frontoparietal attentional networks.

### Valid Retrospective Attentional Modulation Strengthens Target Representations

Extensive behavioral and neural evidence has shown that retro-cues modulate attention in working memory (6,22); however, whether this attention reflects target enhancement or non-target inhibition is still under debate. Recently, an increasing number of studies have pointed out that attentional selection and non-target inhibition are not two sides of the same coin but instead reflect distinct neural mechanisms (32,33). The most vital contribution of the present study is to provide new neurophysiological evidence for an attentional enhancing model of retrospective cues. Behaviorally, we used mixture model to fit behavioral responses and revealed that retrospective attentional cues can enhance target response probability which was consistent with previous studies (34, 35). Furthermore, EEG-based multivariate pattern classification analysis showed that the decoding accuracy of target orientation started to be higher than chance level about 500 ms after the valid retro-cue favoring the target strengthening hypothesis. The decoding accuracy of the target and the behavioral cueing benefits are correlated across individuals. However, the decoding accuracy of non-target orientation after the valid retro-cue kept fluctuating around chance level during the whole retention, not favoring the non-target inhibition hypothesis. The absent observation of non-target inhibition might be due to the memory load in our task was low as previous work found that more non-targets were removed from WM with increasing memory loads (8). Testing non-target inhibition hypothesis with high-load WM tasks might be helpful in future studies.

In neutral condition, the observation that the decoding accuracies of both target and non-target orientations were at chance level, suggesting that both target and non-target representations are held in activity-silent WM depending on short-term synaptic plasticity (36,37). Two similar orientations in the memory space have a close distribution of neuronal coding patterns, which explains why either target or non-target could not be decoded in the neutral condition. The successful decoding of target in valid condition might be due to that the valid retro-cue pings the brain to activate the target representation from the hidden states during retention. Retrospective attentional cues might provide another cluster of neurons involved in the mnemonic representation of the target, thereby amplifying the difference in the mnemonic representation of the target and non-target. Together, these results supported the hypothesis that the target representation was enhanced by the retrospective attentional cue during visual WM.

### Flexible Feature Representations Interact with Retrospective Attention in Visual WM

Previous studies have shown that internal attention can prioritize a subset of mnemonic representations (7,18). Both color and space can be decoded after valid retro-cue, suggesting that these representations of different features are integrated into discrete mnemonic objects (38). Although retrospective cues can enhance the representation of target objects, different features exhibited different temporal dynamics. For color representation during retention and retrieval, a higher decoding accuracy after retro-cue than after probe in valid conditions suggested that effective retrospective attentional cues activate memory representations of task-relevant visual objects through color, reflecting flexible WM representations (39). The encoding of the target space appears to exhibit different neural dynamics compared with the color encoding. Recent studies have found that the micro-saccade moves toward the target location after the center retro-cue (40,41). Our findings claimed that even though the probe is presented in the center of the screen, the task-irrelevant target peripheral location still can be represented in neural activities. A previous micro-saccade study reported that after visiting the location after the retro-cue, unless you did not pay attention during retention, you do not need to go back after the probe (41). Our observations provide further neurophysiological evidence for the findings of eye movements; specifically, although the spatial location is task-irrelevant, retrospective attentional cues can continuously focus attention on the target side, helping to activate and retrieve object-based target representations in WM in advance. Additional, computational models showed that persistent neural activities could be represented by attractor dynamics (42–44). Temporal generation of target space showed that valid retro-cue induced a more stable representation of space than color, indicating that valid retro-cue modulated spatial attention through persistent neural activities. These findings suggested that highly dynamic representations and integration of different features might be good candidates to organize the flexibility of WM representations.

Furthermore, the onset latency of space decoding accuracy is consistent with previous ERP findings in spatial selective attention (45,46). Traditional covert attention contains a physically salient stimulus, which might confuse bottom-up attention capture with top-down attentional selection. Here, we used meaningful color as the retro-cue to isolate top-down control of object held in WM, indicating the object-based attentional modulation in visual WM. Previous studies have claimed that low WM precision is related to spatial binding errors (47,48). Importantly, a negative correlation between space decoding accuracy and response errors revealed that spatial binding of the target object benefited working memory precision. We argued that retrospective spatial attention optimized object-location binding and benefited WM precision. Our observations suggested that object-based representation can be enhanced by retrospective attentional cues for upcoming behavioral performance.

### Mid-frontal Neural Activities Track Trait- and State-level Variability in Attentional Preparatory Processes during Visual WM

We found significant midline prefrontal cortex (PFC) theta ITPC in both conditions and it was correlated with individual behavioral recall errors in each condition, suggesting the critical role of ITPC in representing target in both conditions. These results are consistent with previous results that the midline theta activity in the frontal area is related with cognitive control (49) and that theta phase has been thought to be associated with memory encoding (50). We also found higher ITPC in valid condition, which extended previous observations that theta phase in midline PFC coordinates retrospective attention through brain clocking mechanisms and enhances target representation held in WM.

Corresponding to the higher frontal theta ITPC in valid condition we observed, we also found increased frontal N2 evoked by valid retrospective attentional cue. More importantly, the amplitudes of frontal N2 tracked response speed of WM within individuals, indicating moment-to-moment fluctuations on the attentional control and favoring the ‘readiness to remember’ (R2R) framework (51). This R2R framework explains trait- and state-level variability in mnemonic performance as a function of preparatory attention and goal coding, and their interactions with core mnemonic representations. Our findings suggested theta phase coherence not only the neural marker of behavioral individual differences (theta ITPC), but also of the real-time attentional fluctuations over time (N2), which provides direct neural evidence that both within and between individuals’ variability in attentional preparatory processes influences goal-directed behavior.

We further found that the fronto-central ERP component N600 differed between valid and neutral condition and the difference on N600 amplitudes between two conditions contributed significantly to behavioral retro-cue effects. The positive correlation between retro-cue effect on N600 and retro-cue effect on behavioral error revealed that retro-cue effect on feat binding of the target object benefited working memory precision.

### Organizing Retrospective Attention in Visual WM through Clocking Mechanisms of Theta-alpha Phase-amplitude Coupling in Frontoparietal Connection

The alpha oscillatory activity has been thought to index the deployment of visual attention (23,52) and feature integration (53). Confirming enhanced representation of target space by valid retro-cue in multivariate pattern analysis, we also found larger alpha desynchronization over the parieto-occipital cortex after valid retro-cue, which is consistent with previous studies (21,22,54). Our findings indicated that alpha power shaped the information flow held in WM and modulated the priority of the cued object gated by enhancing spatial attention representation and binding different features.

Previous studies have reported the vital role of frontoparietal network in top-down attentional control (55,56). We found that coupling of theta phases in midline PFC and parietal alpha amplitudes during WM retention was modulated by valid retro-cue, which extends neurophysiological evidence for the engagement of frontoparietal attentional networks in visual WM. This PAC induced by valid retrospective attention was different from many previous studies that found a local PAC in the same brain area (57), because here our PAC result was based on long-range connect between frontal theta phase and parietal alpha amplitudes. Importantly, the increased retro-cue effect on theta-alpha PAC predicted the reduced guessing rate. This long-range frontoparietal connection controlled by retrospective attention supported the fluctuations of top-down information flow in engagement of frontoparietal networks (51), which could guide the goal-directed WM performance.

A previous study found a PAC between frontal theta and parietal alpha in local field potentials when non-human primates performed spatial attention tasks (58). Neural oscillations in the frontoparietal network modulate attention and perceptual sensitivity. Our findings suggested that alpha power in posterior areas might integrate perceptual features into discrete mnemonic objects (38) and theta phase acts as a brain clock to organize retrospective attentional states. A recent opinion proposed that rhythmic attention is a basis of cognitive flexibility (59). Due to the highly flexible representation in WM (39,60), our findings supported that interaction between retrospective attention and flexible mnemonic representation could enhance and prioritize target representation held in WM.

### Conclusions

With a typical retro-cue WM task, we provided convincing neurophysiological evidence supporting object-based attentional strengthening theory. Neural correlates and connections within frontoparietal networks track retro-cue benefits on working memory precision. Retrospective spatial attention optimized object-location binding and benefited WM precision by prioritizing target representations. Our observations provide new insight into training individuals who suffer from memory deficits by modulating attention.

## Methods

### Participants

Thirty-five healthy college students (18 females, age range: 19-28 years) participated in the EEG experiment. No statistical methods were used to previously determine sample sizes, but our sample size is similar to prior studies which applied a similar decoding analysis (61). All of them were right-handed and had normal or corrected-to-normal vision without color blindness. Two participants were removed due to excessive artifacts or ocular movement. Finally, thirty-three participants (17 females, age range: 20-28 years) were involved in further analysis. The current study was approved by the Beijing Normal University Institutional Review Board, and written informed consent was obtained from each participant.

### Stimuli and Procedures

Participants performed a retrospective covert attention task, which required them to select and attend to one of two visual representations maintained in WM. The stimuli were presented on a 21-inch LCD monitor (800 × 600 pixels, 60 Hz) through PsychoToolbox in the MATLAB (The MathWorks Inc., Natick, MA) environment with a viewing distance of 80 cm.

In the encoding stage, two bars surrounded by a ring (size: 4° × 0.32°) were presented at 5° from the left and right of fixation. Bars were randomly allocated to two different colors from blue (RGB: 21, 165, 234), orange (RGB: 234, 74, 21), green (RGB: 133, 194, 18), and purple (RGB: 197, 21, 234), and two different orientations from a set of eight: ±11.25°, ±33.75°, ±56.25°, ±78.75°. The colors were independent across the bar locations and orientations.

Each trial began with an encoding display that contained two to-be-memorized bars (with different colors and orientations) for 200 ms, followed by a delay in which only a central fixation remained on the screen for 1000 ms. Then, a retrospective attentional cue was presented with a color-changed central fixation for 200 ms. In half of the trials, the color was randomly changed to one color of the two bars as a valid cue, indicating which bar needed to be recalled in advance. In the remaining trials, the color was changed to gray as a neutral cue, which provides no attentional benefits for a head start for retrieval. Trials were allocated to valid and neutral conditions randomly within each block. Then, followed by another 1300 ms delay after the retro-cue, a probe bar with a random orientation was displayed in the color of the cued bar (i.e., the target) in the valid-cue trials, while a probe was randomly presented in either color of the two bars in the neutral-cue trials. Participants were instructed to move the mouse and to press the left button to report the orientation of the target as precisely as possible within 4000 ms. After one practice block, the participants completed 20 blocks of 56 trials with a 1-min break between the blocks with EEG recording synchronously.

### Behavioral Modeling

The recall error distribution was first obtained from the angular distance between the reported and the actual orientation for each trial. Then, we fit the response error distribution with the mixture model (62) using the Analogue Report Toolbox in MATLAB. Maximum likelihood estimates of the parameters of the mixture model included the standard deviation (SD) and the probability of the target and uniform responses. SD represents the width of the recall error distribution, indicating an inverse of WM precision. The probability of target and uniform responses represents the probability of correctly reporting the target orientation and guessing a random orientation, respectively.

### EEG Recording and Preprocessing

EEG signals were recorded using a cap with 64 AgCl electrodes in accordance with the international 10-20 system and a SynAmps EEG amplifier while performing the task. Eye movements and blinks were monitored using two electrodes placed 1 cm above and below the left eye for vertical electrooculogram (EOG), and another two electrodes placed at the outer canthus to the eyes for horizontal EOG. Before recording, electrode impedance was maintained below 5 kΩ. Signals were referenced online to the left mastoid, amplified with a bandpass of 0.01-400 Hz, and digitized at a sampling rate of 1000 Hz.

Data preprocessing was performed using the EEGLAB toolbox (63) and custom scripts in MATLAB. EEG signals were first filtered with a bandpass of 0.1–40 Hz and offline referencing to the average of the two mastoids. Segments were then extracted from −2000 ms to 2500 ms relative to the retro-cue onset. Eyeblink and movement artifacts were corrected by independent component analysis (ICA). After ICA, the baseline correction was performed from −200 ms to retro-cue onset in each segment. Then, segments with voltages exceeding ±80 μV at any electrode or exceeding ±50 μV at the horizontal EOG electrode were also excluded. Last, two participants were removed from further analysis if more than 40% of segments were removed due to artifacts or the averaged horizontal EOG activity across the trials exceeded ± 3.2 μV. These procedures removed 2.0% of trials (range: 0–10.5%) from the participants among the final samples, leaving the following number of trials (mean ± std) in each condition: 549±14 for the valid condition, 548±16 for the neutral condition.

### Multivariate Pattern Classification

Multivariate pattern classification was applied to decode the orientations of the target and non-target objects (**Fig 2A**). Each segment for decoding analysis was defined as −200 ms to 2400 ms around the onset of the retro-cue and was down-sampled to 50 Hz to improve the stability of decoding by averaging several adjacent time points (64). Segments were labeled with different target/non-target orientations with 8 labels in total. Several segments with the same labels were averaged into three bins to improve the signal-to-noise ratio. The classifier was based on a linear support vector machine (SVM), and responses in all 60 electrodes serving as features were used to train classifiers through the MATLAB fitcecoc() function at each data point for each participant. The training and testing phases at a given data point were based on different segments. A three-fold cross-validation and ten iterations procedure was applied at each data point to minimize the fortuity caused by the trial assignments and to yield a more stable outcome. Decoding was considered correct only when the target/non-target orientation was determined correctly. Decoding accuracy was computed by comparing the true labels of the target/non-target orientations with the predicted labels, and the chance level was 12.5%.

The color and space of the target objects were also decoded through the same procedure. We calculated the temporal generation of color and space through decoding analysis (65). Specifically, classifiers were trained at a given time sample and evaluated for their ability to generalize to all of the time samples. Therefore, we could obtain a generalization map of decoding accuracy with the training times as the x-axis and the testing times as the y-axis. This approach allowed us to study the temporal dynamics of the color and space representations during WM. A higher than chance level off-diagonal decoding accuracy means that neural representations of color and space remain stable across time (37). Therefore, we quantified the spread time for the decoding accuracy to identify the representation stability of color and space. The specific formulas are as follows:

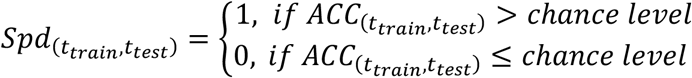

where ACC denotes the decoding accuracy of each training-testing paired time point from the temporal generation map. The chance level for color decoding is 25%, and for space decoding, it is 50%.

Then, to characterize the representation stability, we averaged Spd across the testing temporal dimension by using the following formula to obtain the spread time index over time.

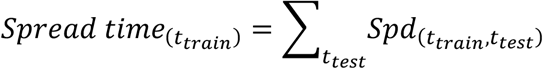

A larger spread time index represents the number of testing time points where the color or space could be decoded at a given training time point, indicating a more stable feature representation.

To examine the significance of the decoding accuracy, segments were arranged to random labels to train and test the control classifiers. Then, a 10000-time cluster-based permutation test was performed across the time and frequencies (66).

### Univariate Analysis

For ERP analysis, each EEG segment was time-locked to the retro-cue onset (–200–2400 ms) and set to a baseline between –200 ms and 0 ms before the onset of retro-cue in each trial. The artifact-free EEG segments were then averaged for valid condition and neutral condition separately. Frontal N600 was averaged between 600 ms and 1300 ms across 9 frontal-central electrodes (F1, Fz, F2, FC1, FCz, FC2, C1, Cz, C2), and averaged frontal N2 was averaged between 320 ms and 360 ms across same 9 frontal-central electrodes. Furthermore, we divided trials into three equal bins according to the behavioral RTs, and then calculated the averaged frontal N2 in each RT bins.

For time-frequency analysis, EEG signals were segmented from –1800 ms to 2400 ms around the onset of retro-cue. To obtain non-phase-locked spectral power, phase-lock ERPs were removed before any spectral measures were calculated to avoid oscillatory activities contaminated by ERPs (67). The instantaneous power and phase at a frequency range of 2–30 Hz in 0.5 Hz steps was estimated by applying a short-time Fourier transform to the Hanning-tapered data. Then, power was baseline-corrected between –1800 ms and –1400 ms (400 ms duration before the encoding onset) and transformed into Log space. Previous studies have reported that alpha power mainly occurs in the posterior cortex; therefore, alpha power (8 – 12 Hz) was estimated in all parieto-occipital electrodes (P3/4, PO3/4, PO5/6, P7/8, PO7/8, O1/2, Pz, Oz). Averaged posterior alpha power (8 – 12 Hz) in both valid and neutral conditions for statistic were then calculated between 300 and 600 ms.

The inter-trial phase coherence (ITPC) was calculated across trials through newtimef function in EEGLAB, reflecting the consistency of phase values across trials within each electrode. The ITPC is stimulus-locked and independent of amplitude changes. The value of ITPC is between 0 and 1. A value closer to 0 indicates a lower phase synchronization across trials, and a value closer to 1 represents a higher phase synchronization across trials. Here, ITPC in theta band (3 – 7 Hz) was calculated in the midline frontal electrode (Fz) and averaged between 100 ms and 700 ms to examine the difference between valid and neutral conditions.

For phase-amplitude-coupling (PAC) analysis, we first extracted alpha amplitudes and theta phases by applying a Hilbert transform (hilbert.m) on filtered EEG signals. To acquire a stable estimation of PAC, we maximized the measurement window to the whole retention (0 – 1500 ms after retro-cue onset). Theta phases were calculated in the midline frontal electrode (Fz) and alpha amplitudes were calculated in all electrodes to examine connections between mid-frontal cortex and the whole brain. Then, the modulation index was calculated to assess PAC in both valid and neutral conditions (68). A larger PAC modulation index means stronger coupling between two frequencies. The PAC index were then averaged across all parieto-occipital electrodes (same with electrodes used to estimate alpha power) to examine the difference between valid and neutral conditions.

## Acknowledgments

The present research was supported by the National Natural Science Foundation of China Grant Nos. 31871099 (to YS), and the National Defense Basic Scientific Research Program of China Grant No. 2018110B011 (to YS).

## Data and Code Availability Statement

Data and codes supporting the findings of this study will be available upon reasonable request to corresponding authors.

